# Estimation of auditory steady-state responses based on the averaging of independent EEG epochs

**DOI:** 10.1101/438010

**Authors:** Pavel Prado-Gutierrez, Eduardo Martínez-Montes, Alejandro Weinstein, Matías Zañartu

## Abstract

The amplitude of the auditory steady-state responses (ASSRs) generated in the brainstem exponentially decreases over the averaging of subsequent EEG epochs. This behavior is partially due to the adaptation of the auditory response to the continuous and monotonous stimulation. We analyzed the potential clinical relevance of the ASSR adaptation. Specifically, we compare the ASSR amplitude computed in two conditions: (1) when the auditory responses -embedded in the EEG epochs that are averaged in the estimation procedure- are influenced by the previous stimulation; and (2) when they are independent of the previous stimulation. ASSR were elicited in eight anesthetized adult rats by 8-kHz tones, modulated in amplitude at 115 Hz. ASSR amplitudes were computed using three averaging methods (standard, weighted and sorted averaging). We evaluated the ASSR amplitude as a function of sub-set of epochs selected for the averaging and the improvement in the ASSR detection resulting from averaging independent epochs. Due to adaptation, the ASSR amplitude computed by averaging dependent EEG epochs relied upon the averaging method. Lower ASSR amplitudes were obtained as EEG segments containing unadapted responses were systematically excluded from the averaging. In the absence of EEG artifacts, the ASSR amplitudes did not depend on the averaging method when they were computed from independent EEG epochs. The amplitude of independent ASSRs were up to 35% higher than those obtained by processing dependent EEG segments. Extracting the ASSR amplitude from independent epochs halved the number of EEG segments needed to be averaged to achieve the maximum detection rate of the response. Acquisition paradigm based on a discrete acoustic stimulation (in which segments of AM-sounds of several seconds in length are presented after a given inter stimulus interval), in combination with appropriated averaging methods might increase the accuracy of audiological tests based on ASSRs.

## Introduction

Auditory steady-Sastate responses (ASSRs) are brain oscillations locked to the periodic properties of acoustic stimuli (Picton et al. 2003; John and Purcell 2008). Audiological tests based on the acquisition of ASSR are useful for estimating the hearing sensitivity, mainly because multiple hearing frequencies can be simultaneously assessed, and the auditory response can be objectively detected using statistical tests (Savio et al. 2001; Luts et al. 2004; Valdes et al. 1997; Wilding et al. 2012; de Resende et al. 2015).

Typically, ASSR are elicited by the continuous presentation of amplitude modulated (AM) tones. The extraction of the auditory response from the measured signal essentially relies on averaging epochs of the EEG, time-locked to the stimulus (Dawson 1954). Such a manipulation assumes that the EEG signal is a linear superposition of the highly stereotyped, time-invariant response, and the ongoing background noise (Glaser and Ruchkin 1976). However, evidences obtained in several sensory pathways suggest that the evoked potential amplitude might not be steady but decreases exponentially due to the serial and regular stimulation (Pereira et al. 2014; Andrade et al. 2015; Custead et al. 2015). Such effect has been defined as evoked potential adaptation.

Evidences supporting the adaptation of auditory evoked potentials (AEP) has been provided by analyzing the effect of the stimulation rate on the amplitude and latency of transient responses. Those studies show that, as the presentation rate of acoustic stimuli increases, the amplitude of AEPs obtained in both humans and rodents decline (studies in humans: Ritter et al. 1968; Rosburg et al. 2010; Zacharias et al. 2012; studies in rodents: Knight et al. 1985; Sambeth et al. 2004; Budd et al. 2013; Duque et al. 2018). When the time-course of transient AEPs amplitude has been analyzed, it has been observed that the asymptotic amplitude of the response is preceded by an initial stage, in which the amplitude decreases over several stimulations (Ritter et al. 1968; Pereira et al. 2014; Paiva et al. 2016; Rosburg and Sörös 2016).

Traditionally, it has been argued that the ASSRs primarily result from the linear superposition of transient AEPs elicited by the high presentation rate of acoustic stimuli (Galambos et al. 1981; Tan et al. 2017). Nevertheless, unlike the suppression of transient AEPs induced by the stimulus repetition, the adaptation of ASSRs has received relatively little attention. Although several studies have analyzed the time course of the ASSR amplitude (John and Picton 2000; John et al. 2001; Torres-Fortuny et al. 2011), they have focused mainly on describing the variations in amplitude resulting from the time-domain averaging of sequentially acquired EEG epochs, thus, assuming that the amplitude of the response is the same over time. We have noticed that this averaging procedure does not allow to discriminate between methodological and physiological related variations in the amplitude of the evoked response (Prado-Gutierrez et al. 2015). For testing the adaptation of the evoked potential, it is necessary to obtain a reliable response at any given epoch, independently of the electrical activity of the preceding EEG segments. In that scenario, similar ASSR amplitudes estimated independently of the preceding electrical activity strongly support the strict stationary behavior of the ASSR. Alternatively, a decrease of amplitude along independent epochs accounts for the adaptation of the response.

Using the methodology proposed by Ritter et al. (1968) for quantifying the adaptation of cortical AEP, we have demonstrated that the amplitude of ASSRs generated in the brainstem decrease exponentially due to the sustained presentation of AM sounds (Prado-Gutierrez et al. 2015). Such a behavior might reflect the loss of novelty of the sensory input, increasing the sensitivity to relevant fluctuations in the acoustic environment (Ritter et al. 1968, Malmierca et al. 2009).

The adaptation of ASSR might have implications in the clinical practice, especially when recording protocols are based on averaging a reduced number of sequentially acquired epochs (Luts et al. 2008; Choi et al. 2011; Wilding et al. 2012). In such a practical situation, the ASSR computed after the completion of the averaging might be strongly influenced by the adaptation of the response. This is important considering that the ASSR amplitude estimated at the end of the recording is used to judge the significance (statistical detection) of the auditory responses.

Possible shortcomings in the computation of ASSRs resulting from adaptation might be overcome by implementing stimulation protocols which prevent the suppression of the ASSR amplitude over time. Additionally, a better estimation of the response might result from the implementation of averaging procedures that attenuate the effect of motion and muscular artifacts, i.e., using weighted and sorted averaging methods (Hoke et al. 1984; Elberling and Wahlgreen 1985; Lütkenhöner et al. 1985). Weighted averaging involves normalizing the voltage samples of each individual EEG epoch by an estimate of the amplitude variability, e.g., weighting the data samples by the inverse of either the variance, or the standard deviation of the voltage amplitude (Hoke et al., 1984). Sorted averaging comprises the rearrangement of epochs as a function of the voltage variability, averaging only those epochs which contribute to increasing the accuracy of the response estimation. The latter is typically achieved by sorting the epochs in an ascending order of their root mean square (RMS) and averaging first those epochs with low RMS, as they contribute to increasing the SNR of the signal (Mühler and von Specht 1999; Rahne et al. 2008). Both weighted and sorted averaging have been applied to the analysis of ASSR. Weighted averaging is already available in commercial ASSR systems, and it is commonly used for research purposes (John et al. 2001, Wilding et al. 2012; Wilson et al. 2016). Sorted averaging has only been tested experimentally (Rahne et al. 2013), probably due to the relatively high computational cost of storing and sorting a large number of epochs during the online estimation of ASSRs.

In this study, we analyzed possible biases in the computation of the ASSR amplitude resulting from adaptation. More specifically, we compared the ASSR amplitudes obtained in two conditions: (1) when the auditory responses -embedded in the EEG epochs that are averaged in the estimation procedure- are influenced by the previous stimulation; and (2) when they are independent of the previous stimulation. We hypothesize that the detection of ASSR will significantly improve when the ASSR is computed by averaging EEG epochs containing only unadapted auditory responses. We tested this hypothesis by thoroughly characterizing the brainstem ASSR of rats, computed with standard, weighted, and sorted averaging methods. Using the Hotelling T2 test for the detection of the response (Valdes et. al 1997), the improvement in the ASSR detection resulting from averaging independent epochs was also evaluated. The results are discussed based on a comparison with existing paradigms designed to optimize the detection of ASSR.

## Materials and Methods

### Experimental subjects

Auditory responses were obtained from 8 adult Wistar rats. Animals were housed in a standard bio-clean animal room under a 12-h light-dark cycle at 22-24°C, with free access to food and tap water. To perform the recordings, animals were anesthetized with ketamine (75.0 mg/kg, ip) and diazepam (5.0 mg/kg, ip). Supplemental doses of anesthesia were administered during the experiment at a level sufficient to maintain the animal in an areflexic state. Atropine sulfate (0.06 mg/kg; im) was administered to decrease the mucosal secretions. Body temperature was maintained at 37.0±0.1°C by a body temperature control system (Bioseb, model LE-6400). The present study was performed under approval of the Animal Research and Ethics Committee of the Cuban Neuroscience Center, conformed to the guidelines of the National Center for Animal Breeding of Cuba.

### Acoustic stimuli

Continuous tones of 8 kHz sinusoidally-modulated in amplitude (95% depth) at 115 Hz were generated using the ASSR software module (Perez-Avalo et al. 2001) of the AUDIX system (Havana, Cuba) and presented monaurally at 50 dB SPL, via an ER 3A Etymotic Research insert earphone. Custom-fitted ear molds were used to replace the original foam to permit the earphone to be coupled to the rat’s ear. Acoustic stimuli of 8 kHz have been previously used to study the ASSR in rats (e.g., Perez-Alcazar et al. 2008; Prado-Gutierrez et al. 2012; Venkataraman and Bartlett 2014), since this frequency corresponds to a peak in the spectral hearing sensitivity of the specie (Kelly and Masterton 1977; Heffern et al. 1980). The acoustic levels are referred to a Brüel & Kjær artificial ear (type 4152). Calibration was performed using a Brüel & Kjær 2250 sound level meter (Brüel & Kjær 4144 microphone).

### Recordings

Electrophysiological responses were recorded differentially using stainless-steel needle electrodes inserted subdermally (vertex positive; neck negative; thorax reference). Recordings were amplified with gain 1.2×10^4^ and band-pass filtered −cutoff frequencies of 10 and 300 Hz. Output of the filter was digitized at 16 bit of resolution and sampled at 920 Hz. Segments with peaks of electrical oscillations exceeding 50 mV were rejected online. Typically, less than five segments per recording were rejected and they were randomly distributed across epochs. Data acquisition continued until completing 60 artifact-free epochs of 4.45 s (4096 time-points each). Thirty recordings were acquired from each animal. During the experimental session, every recording was preceded by a no-stimulation period of ten minutes.

### Pre-averaging modifications of epochs

Processing of the data was performed using in-house Matlab codes (MathWorks, USA). The 60 sequential epochs of the 30 recordings were re-arranged offline into a data matrix of 30 rows and 60 columns -one matrix per animal (Fig 1, left panel). From this dataset we created two other modified data matrices, containing weighted and sorted epochs, respectively (Fig. 1A, middle and right panels). Both manipulations are based on assuming that the sample variability of the epoch reflects only the contribution from noise, so that noisy epochs will have higher amplitude variability (Hoke et al. 1984; Mühler and von Specht 1999).

**Fig 1.**
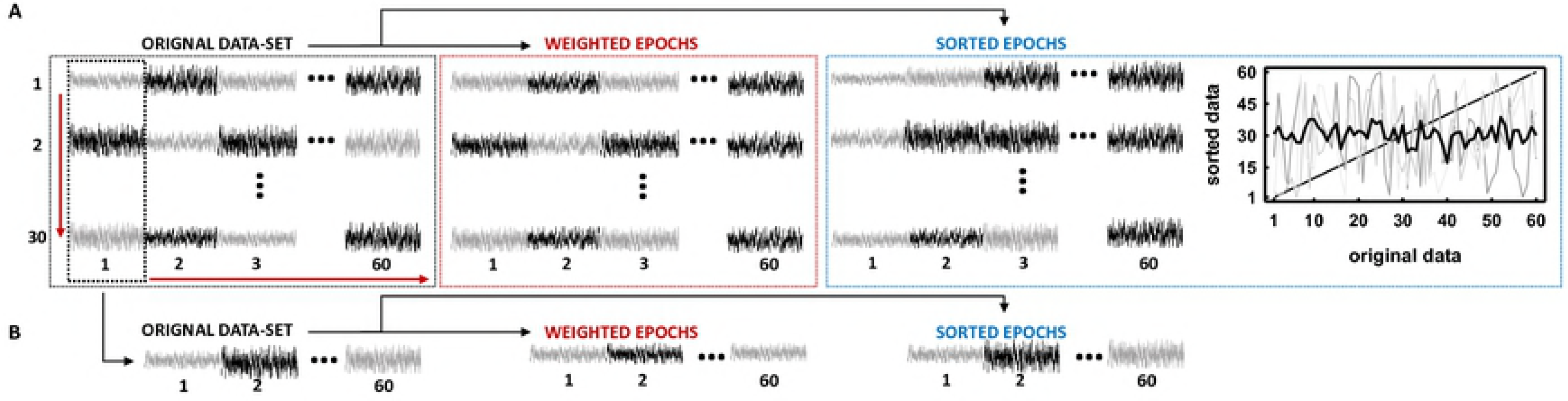
Diagram illustrating the arrangement of measured EEG epochs as a data matrix with 30 rows (independent recordings) and 60 columns (dependent epochs); and the averaging methods used for the estimation of ASSR. A) Modifications of the original data-set for performing weighted and sorting averaging. An example of sorting is presented in the right-most panel, where grey plots represent the location of epochs of individual recordings before (x-axis) and after (y-axis) the sorting. The black plot represents the mean location through the 30 recordings. Red arrows represent the classical averaging of sequential epochs within a recording (horizontal arrow) and the averaging of independent epochs corresponding to the same time window in each of the 30 recordings (vertical arrow). B) Epochs corresponding to the first-time window in the different recordings were concatenated to obtain a new synthetic recording formed by independent epochs, to estimate unadapted ASSR amplitudes.

Weighted epochs were obtained by dividing each voltage sample by the amplitude variance of the epoch they belong to, so that variance was used as a measure of variability and weighting factor (John et al. 2001). Pertinently, we normalized the weights by their average across all epochs in order to make the ASSR amplitudes obtained with three averaging procedures comparable. In the sorting procedure, epochs were rearranged following an ascending order of root-mean-square (RMS). This parameter has been used before as sorting factor in the detection of ASSRs, as it is assumed to be proportional to the level of background noise (Rahne et al. 2013). In our case, the sorting procedure was performed separately for each recording, which implied that epochs corresponding to the same time window in different rows of the original matrix, might appear in different time windows of the ordered dataset (Fig.1 right panel).

### Adaptation of ASSRs

The adaptation of the ASSR was analyzed as described in Prado-Gutierrez et al. (2015). In summary, the averaging of epochs in the data matrices was performed along columns (Fig. 1A). This was performed for each dataset (original, weighted and sorted epochs). it is important to recall that sorting was performed according to the RMS values along each row, but the averaging was carried out along columns, thus adding together epochs that do not correspond to the same time window in the original dataset.

The amplitude of the ASSR was computed once for each group of epochs, at the end of the averaging, using the fast Fourier transform (FFT). Such a manipulation allowed us to explore the time evolution of the ASSR amplitude along a recording (a row in the data matrices) without the influence of noise. The spectral amplitude obtained at 115 Hz (frequency of the amplitude modulations of the acoustic stimuli) was considered as the amplitude of the ASSR. The amplitude of the 30 spectral components at each side of the frequency of the response were vector averaged to calculate the residual noise level (RNL). Those spectral components corresponded to the frequency range between 111.8 Hz and 118.2 Hz. The ASSR amplitudes estimated along column can be considered independent of one another, since averaging did not involve different epochs of the same recording. Hereinafter, they will be called independent responses.

Both the independent ASSR amplitude and the RNL were plotted as a function of the position of the epochs within recordings, i.e. the number of the column in the data matrix (see Fig. 1A). One-way ANOVAs (p<0.05) and the corresponding post-hoc analyses (Tukey test, p<0.05) were performed to analyze the changes in the amplitude and the RNL of the independent responses. Since the adaptive behavior of the ASSR described in Prado-Gutierrez et al. (2015) was evident during the first 30-s of stimulation, the statistical analysis of adaptation performed in this study including only the first 10 independent responses. The same reasoning applies for the rest of the statistical analyses described below.

### Dependent ASSR amplitudes

For analyzing the dynamics of the ASSR amplitude during the averaging procedure, epochs corresponding to the same recording were sequentially averaged in the time-domain using standard, weighted and sorted averaging. The direction of the averaging is represented in the middle panel (Fig. 1A). The ASSR amplitude was estimated after including each additional EEG epoch in the averaging. We will refer to this procedure as averaging of “dependent epochs” and correspondingly, estimation of “dependent ASSR amplitudes”, as subsequent values were estimated by using the same epochs but one.

Furthermore, we analyzed whether the time evolution of dependent ASSR amplitudes varied as different pools of epochs within the same recording were averaged. Specifically, we excluded from the averaging the first 1, 2, 4, 8 and 16 epochs of the recording, in order to evaluate the specific influence of these first epochs in the estimated ASSR amplitudes. The rejection of epochs was performed after the modifications associated with weighted and sorted averaging.

A three-way ANOVA (p<0.05) and the corresponding post-hoc analyses (Tukey test, p<0.05) was conducted to compare the ASSR amplitudes obtained at the end of the averaging as a function of the averaging method, the number of averaged epochs, and the number of epochs excluded from the averaging.

### Independent vs. dependent ASSR amplitudes

The aim of this study is testing whether the averaging of independent EEG epochs results in higher ASSR amplitudes than those obtained by averaging dependent EEG segments. To this end, the first epochs in each recording (first column in the original data-set) were concatenated to form a synthetic recording of epochs that are independent of one another (Fig. 1B). As mentioned before, the auditory response embedded in any of those epochs are not affected by the preceding responses, since recordings were obtained after ten minutes of resting. Weighting and sorting of epochs were performed after the concatenation procedure (Fig. 1B middle and right panels) and epochs were averaged in the time-domain. The ASSR amplitude was estimated after including each additional EEG epoch in the averaging, which allows to find the variations in the response amplitude as a function of the number of averaged epochs. This was then contrasted with the ASSR amplitudes computed averaging subsequent epochs within the original recordings. A three-way ANOVA (p<0.05) was performed to compare the ASSR amplitudes, using the type of epochs (dependent vs. independent), the averaging method (standard vs. weighted vs. sorted) and the number of averaged epochs as factors.

### Detection rates

The ASSR amplitudes and the RNL were compared by using the Hotelling's T2 multivariate test implemented in the AUDIX system (Valdes et al. 1997), which considers both the amplitude and phase of the oscillations. The statistical test was performed after averaging each additional EEG epoch, both in the case of dependent and independent ASSR responses and using the three different averaging methods. ASSRs were considered as detected when the response amplitude was significantly higher than the RNL. Detection rates were computed at each time window as the fraction of animals where the response was statistically significant.

## Results

### ASSR adaptation

Fig 2 (upper panels) illustrates the ASSR amplitudes calculated for independent epochs. The adaptation of the ASSR was evident by analyzing the original and weighted data-sets. In both cases, the independent ASSR amplitudes decreased over the first four EEG segments and remained steady afterward (F=17.30, p<0.05 and F=14.66, p<0.05 for original and weighted epochs, respectively). However, the adaptive behavior of the ASSR was not detected in the sorted dataset (F=0.88, p>0.05). As expected, the RNL of the signals did not vary as a function of the averaging method (Fig 2, lower panels) (F=0.89, p>0.05, F=0.93, p>0.05 and F=0.25, p>0.05 for original, weighted and sorted epochs, respectively).

**Fig 2.**
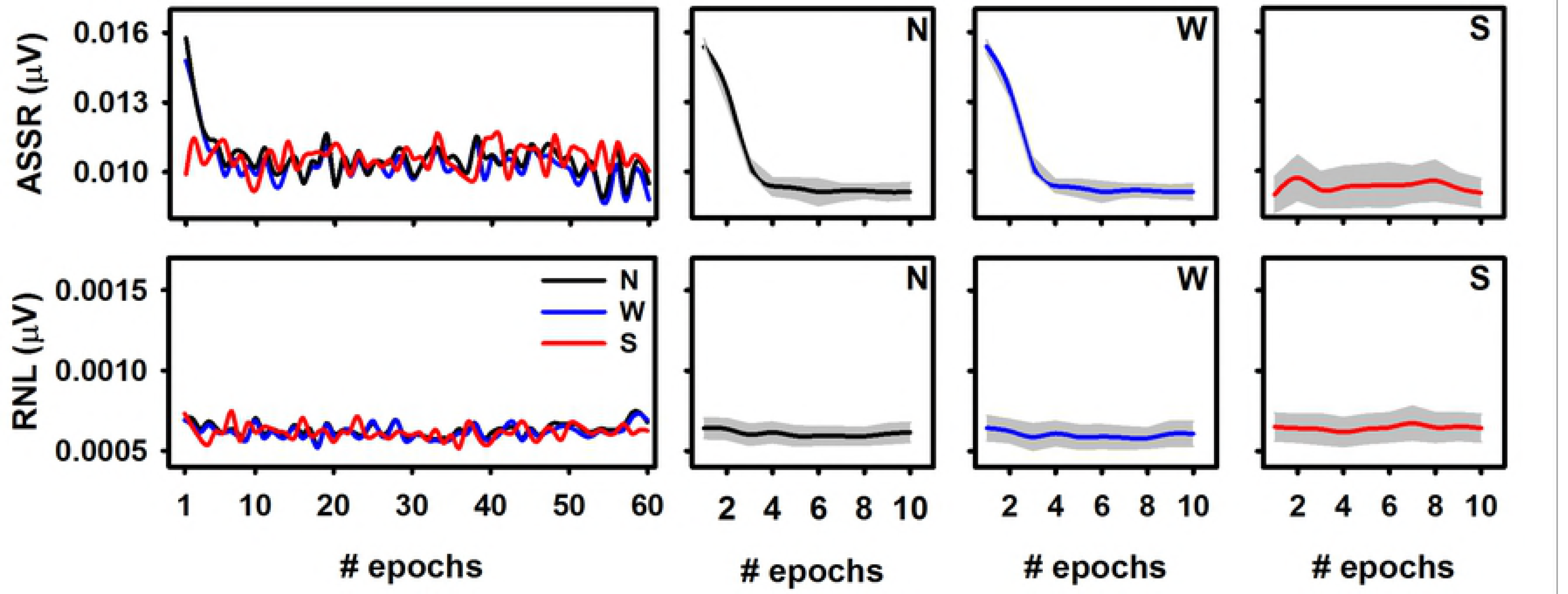
Independent ASSR amplitudes (upper panels) and the corresponding residual noise level (RNL, inferior panels) plotted as a function of the number of epochs. Traces corresponding to a typical individual are displayed in the left panels. Means and standard deviations (N=8) are plotted in the rest of the panels for original (N), weighted (W) and sorted (S) data-sets.

### Evolution of the dependent ASSR amplitudes

The time course of the ASSR obtained during the standard sequential averaging of epochs within a recording (dependent epochs) was characterized by a progressive decrease of amplitude, which was mainly evident during the averaging of the first EEG segments (Fig 3, left upper panel). Such a behavior depended on the pull of epochs selected for the response estimation. In other words, when the response evaluation was initiated after a given number of epochs were acquired, lower initial ASSR amplitudes were computed as larger sub-set of epochs acquired at the beginning of the recording were subsequently excluded from the averaging. Consequently, lower decreases of dependent ASSR amplitude were systematically obtained as the beginning of the response evaluation was delayed, i.e., as more of the epochs acquired at the beginning of the recording were systematically excluded from the averaging. In fact, the ASSR amplitude decreased only slightly during the averaging procedure, when the first 16 epochs of the recording were ruled out. Similar dynamics resulted from applying weighted averaging (Fig 3, middle upper panel). Nevertheless, when the ASSRs were estimated using sorted-averaging, the response amplitude was relatively constant as the number of averaged epochs increased (Fig 3, right upper panel). Furthermore, the behavior of the ASSR amplitude during the sorted averaging procedure did not vary as a function of the number of epochs ruled out before the beginning of the response evaluation.

**Fig 3.**
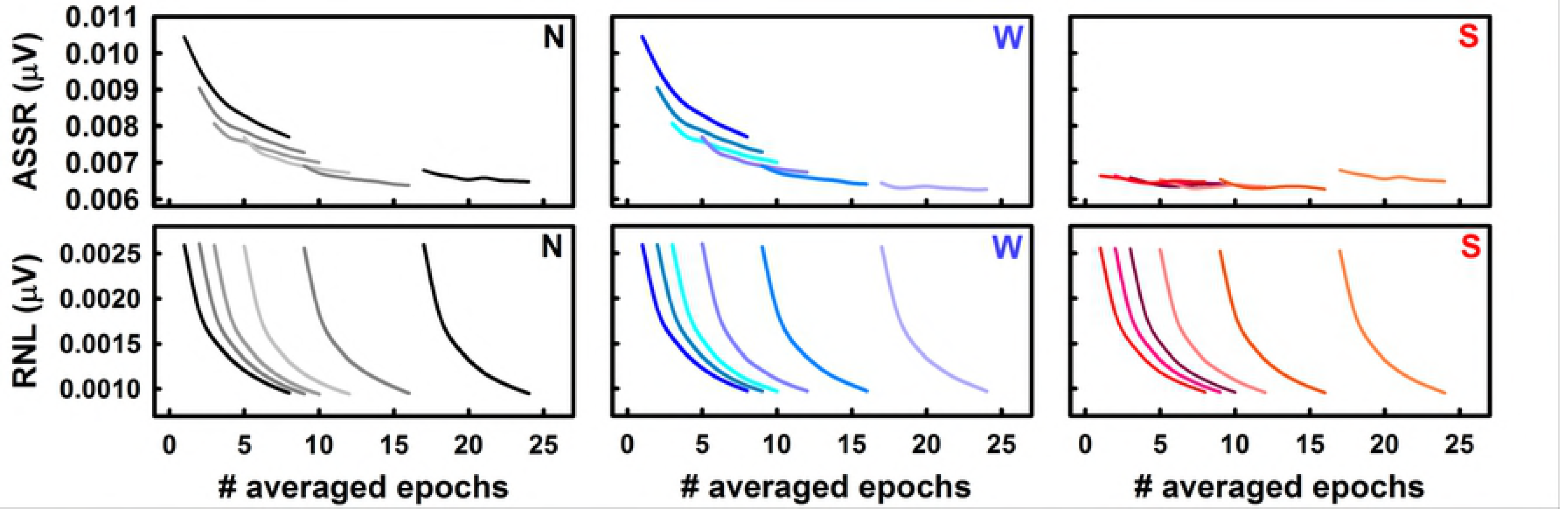
Evolution of the dependent ASSR amplitude during the standard (N), weighted (W) and sorted (S) averaging of eight EEG epochs. Traces represents the mean amplitudes of 30 recordings obtained in a representative individual. The response estimation started after rejecting a given sub-set of epochs acquired at the beginning of the recording (0, 1, 2, 4, 8 or 16 epochs).

Consequently, the ASSR amplitude obtained at the end of the averaging depended on both the averaging method and the size of the data-subset excluded from the analysis (F=5.66, p<0.05 and F=3.20, p<0.05 for the effects of the averaging methods and the number of rejected epochs, respectively). When using the standard and weighted averaging methods, the ASSR amplitude computed by averaging only two EEG epochs significantly decreased as the number of excluded epochs increased up to eight (Fig 4, left panel). Further increases in the number of rejected epochs did not have a significant effect on the response amplitude. Once again, a different behavior resulted from applying sorted-averaging. When this method was implemented, the ASSR amplitude did not significantly vary as a function of the number of epochs ruled out for the averaging. Furthermore, when up to four epochs were excluded from the averaging, the sorting of epochs resulted in lower ASSR amplitudes as compared to those obtained with the standard and weighted averaging protocols. The same trends were obtained when the ASSR amplitude was estimated by averaging four and eight epochs (Fig 4, middle and right panels).

**Fig 4.**
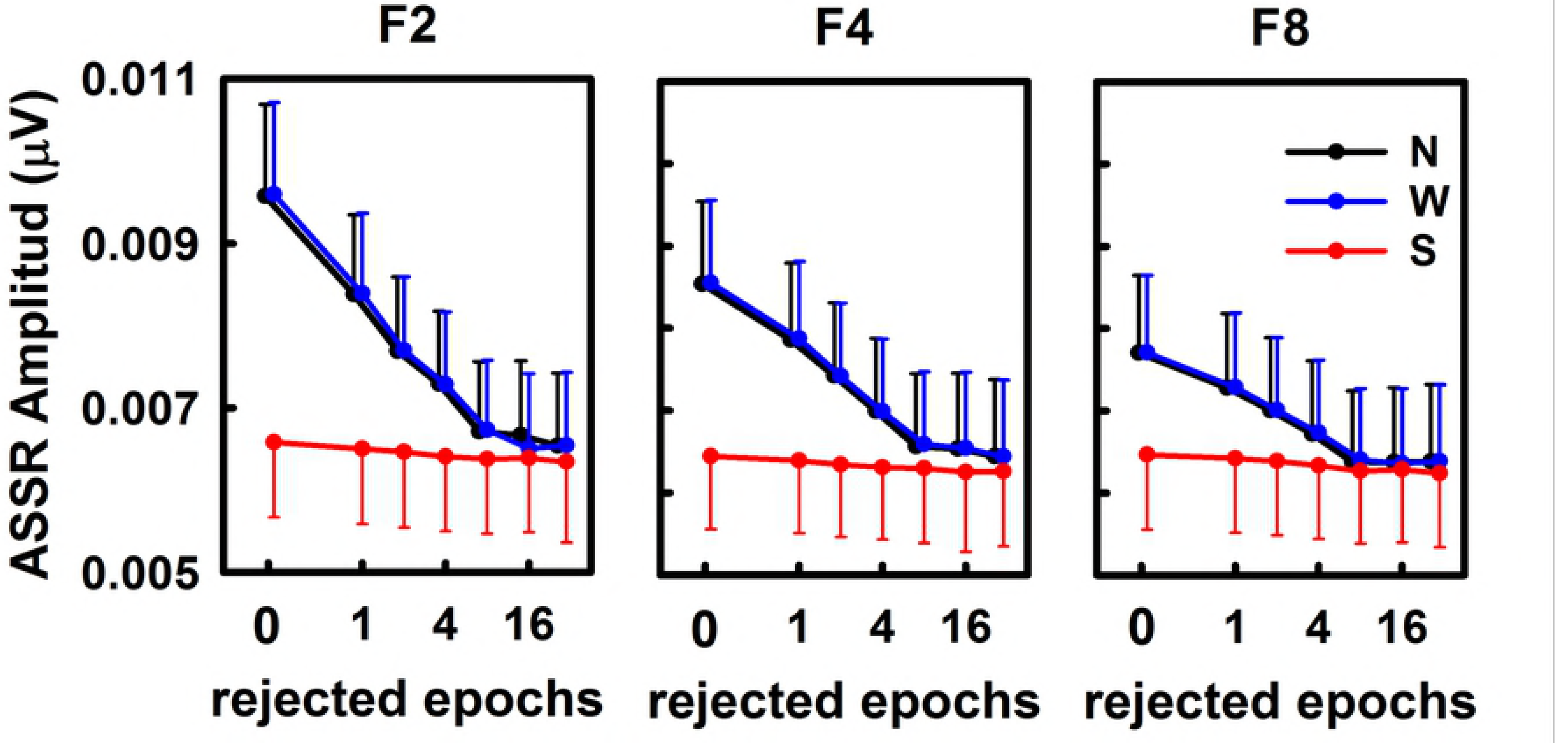
Estimation of dependent ASSR amplitudes as a function of the size of the sub-set of epochs acquired at the beginning of the recording that were excluded from the averaging. F2, F4 and F8 represent the fixed number of averaged epochs (2, 4 and 8 epochs, respectively). Averaging protocols are also represented (N: standard, W: weighted, S: sorted). Plots represents the mean (symbol) ± standard deviation (vertical bar) across eight individuals.

The changes in the ASSR described above could not be explained by the behavior of the RNL during the averaging procedure. Increasing the size of the data subset excluded from the averaging did not have any effect on the behavior of the RNL (Fig 3, bottom panels). Furthermore, the dynamics of the RNL did not change as a function of the averaging methods The RNL estimated with every protocol exponentially decreased as the averaging of epochs within the recordings was performed.

### Independent vs. dependent ASSR amplitudes

Fig 5. illustrates a comparison of the ASSR amplitudes computed during the averaging of dependent epochs within a recording and those obtained by averaging independent epochs containing unadapted auditory responses. By way of reminder, independent responses were obtained by concatenating the first epoch of the 30 recordings acquired in each animal (Fig.1B).

**Fig 5.**
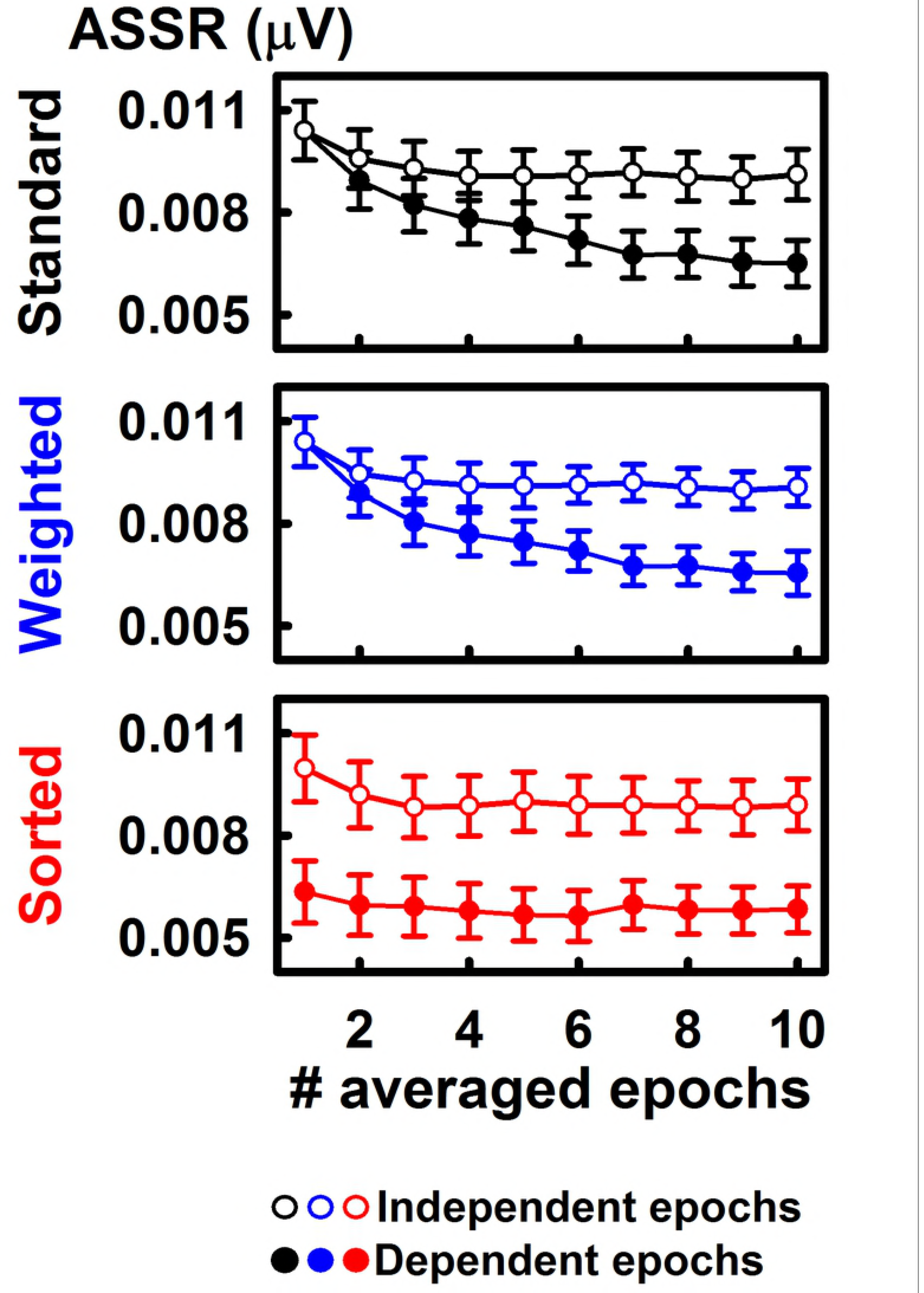
ASSR amplitudes computed by averaging dependent and independent EEG epochs. The dynamics resulting from applying standard, weighted and sorted averaging are represented. Plots represent the mean ± standard error (N=8).

Unlike the time evolution of the ASSR amplitude obtained during the averaging of dependent EEG epochs, the independent ASSR amplitude did not varied as a function of the averaging method (Fig. 5). Furthermore, averaging independent epochs consistently resulted in higher ASSR amplitudes than those evoked by averaging dependent EEG segments (F=6.83; p<0.05). The benefit of computing the independent ASSR amplitudes using the standard and weighted averaging was evident after acquiring at least six EEG sepochs (F=11.75; p<0.05). Using sorted averaging, the independent ASSR amplitudes were significantly higher than those computed for dependent EEG segments, even from the first epoch. When the auditory response was evaluated after averaging 10 epochs, the ASSR increased by 28.3%, 27,8% and 34.5% when they were estimated with standard, weighted and sorted averaging, respectively.

The changes in the ASSR resulting from averaging independent instead of dependent EEG epochs were reflected in the detection rate of the auditory response. As expected, when any of the three averaging methods were used, the detection rates of the ASSR increased as more dependent or independent epochs were averaged (Fig 6). When the ASSR amplitude was computed by standard and weighted averaging, the initial detection rates of dependent and independent responses were similar (Fig 6 A and B). Nevertheless, the initial detection rate obtained by the sorted averaging of independent epochs was 45 % higher than that computed by averaging sorted dependent EEG segments (Fig 6 C). More importantly, extracting the ASSR amplitude from independent epochs halved the number of EEG segments needed to be averaged to achieve the maximum detection rate of the response.

**Fig 6.**
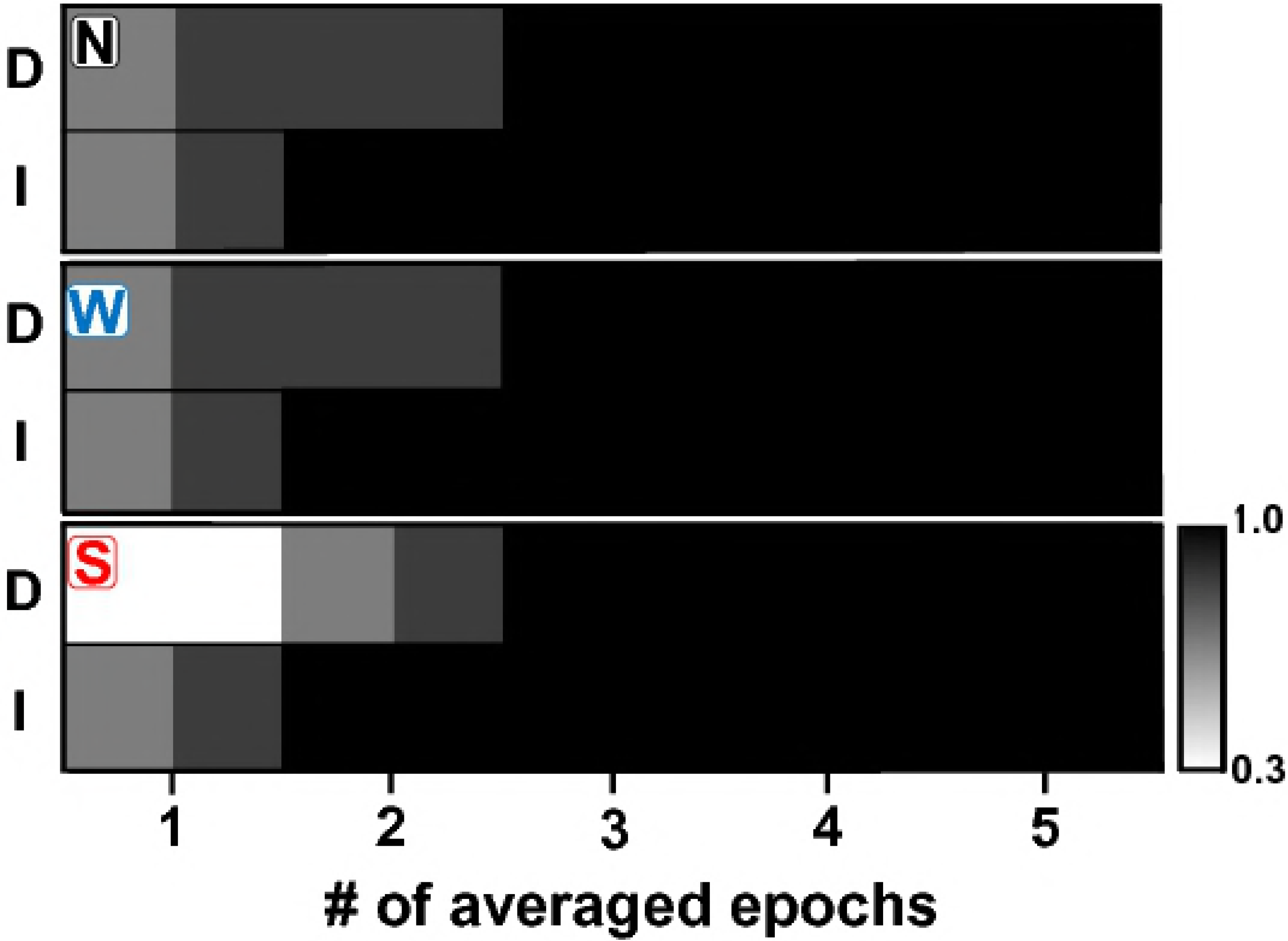
Detection rates of ASSR elicited by averaging dependent (D) and independent (I) epochs, as a function of the number of averaged epochs. Detections rates resulting from standard (N), weighted (W) and sorted (S) averaging are represented in the upper, middle and lower panels, respectively.

## Discussion

In this work we tested the theoretical principles of an acquisition paradigm for reducing the adaptation of the ASSR. Our results demonstrate that, in absent of EEG artefacts, the ASSR amplitude estimated by processing dependent EEG epochs vary as a function of the averaging method used in the estimation procedure. The effect of the averaging method is not evident when the ASSR amplitudes are computed by averaging independent EEG epochs. More importantly, our results demonstrate that averaging independent EEG epochs result in significantly higher ASSR amplitudes than those obtained by averaging dependent EEG segments. Consequently, averaging independent instead of dependent EEG epoch significantly improves the detection of ASSRs.

### ASSR adaptation

The exponential decrease of independent ASSR amplitudes described in Fig 2 (upper panels) replicates our previous findings and supports the notion that the ASSR adapts to the continuous presentation of acoustic stimuli (Prado-Gutierrez et al. 2015). These results are in accordance with studies describing the adaptation of the transient AEP in both humans and animal models (Ritter et al. 1968; Öhman and Lader 1972; Bourbon et al. 1987; Zhang et al. 2009; Pereira et al. 2014; Paiva et al. 2016; Rosburg and Sörös 2016, Duque el al. 2018).

In this study we also provide a more precise description of the ASSR adaptation by using smaller FFT windows that than applied in our previous study. Noteworthy, we founded new evidences of the ASSR adaptation by analyzing the independent ASSR amplitudes of weighted and sorted epochs. On one hand, the ASSR adaptation is evident after the weighting procedure. This result is explained by the fact that we used the variance of each epoch as the weighting factor and that this parameter does not depend on the mean amplitude value of the epoch. Due to the small amplitude of the auditory response relative to the background noise, the variance of the epoch mainly reflects the variance of the noise. As the level of noise was found to be the same in both epochs containing unadapted and adapted ASSR (Fig 2 and Prado-Gutierrez et al. 2015), the weighting did not affect the amplitude of the ASSR embedded in all epochs.

On the other hand, relatively constant amplitudes of the independent ASSR were obtained after sorting the epochs in an ascending order of RMS within the recordings (Fig. 2). This was a consequence of the small contribution of the auditory response to the RMS of single epochs when compared with the contribution of the background noise, even in those epochs containing unadapted ASSR. Therefore, after sorting, epochs containing unadapted ASSR will not be generally placed at the beginning but in any other location of the recording. Consequently, any given column in the sorted data-set (Fig 1) will be mainly composed by epochs containing adapted ASSR. Since independent ASSR were computed by averaging through the columns in the sorted data matrices, similar independent ASSR amplitudes are expected due to the relatively equal contribution of unadapted epochs to every time window.

From a phenomenological perspective, the behavior of the ASSR described in this study meet the principal criteria defined by Thompson and Spencer (1966) for adaptation: the asymptotic decrease of the response amplitude. As recently suggested by Duque et al (2018) analyzing auditory brainstem responses (ABR) of anaesthetized rodents, the adaptation of scalp recorded AEP reveals the adaptation of specific neural populations in the auditory pathway. However, in addition to adaptation, other physiological processes such as refractoriness might also contribute to the dynamics of the ASSR. As suggested by the experimental results and the theoretical model presented by Zacharias et al. (2012), refractoriness might play a relevant role at ranges of time shorter than 5 s. Therefore, it can be speculated that the balanced activation of a sub-pool of neurons which are refractory to the stimulation and another composed by neurons which are in a recovery-after-refractoriness stage, might contribute to explain the asymptotic amplitude of the ASSR.

### Cortical vs. brainstem ASSR adaptation

A recent study, using the methodology implemented in Prado-Gutierrez (2015) for quantifying the ASSR adaptation, reported a significant but very weak decrease in the amplitude of the human 40-Hz ASSR over time, concluding that the 40-Hz ASSR does not adapt to the continuous stimulation (Van Eeckhoutte et al. 2018). Those authors accounted for the discrepancies with our results based on differences in the ASSR neural generators (cortical versus brainstem) and differences between species (humans versus rats).

The reduced adaptation of the cortical ASSR can be explained as part of the gradient in the levels of neural adaptation existing from the auditory periphery to the cortex (Loquet et al. 2004; Meyer et al. 2007). Such a gradient is reflected in the different adaptation pattern of the human ABR with respect to that of the auditory middle latency response (MLR) (Özdamar and Bohórquez 2007; Özdamar et al. 2007). It is also evident when analyzing the sensitivity to the inter-stimulus interval (ISI) of the earlier relative to the late components of the AEP in rats (Budd et al. 2013).

Nevertheless, it is important to note that the anatomical organization and the physiological mechanisms of the auditory system are very consistent across mammals. These homologies are reflected in several properties of AEP recorded from humans and rodents (Boutros et al. 1997; Shaw 1988; de Bruin et al. 2001). Remarkably, similarities between humans and rodents have been well documented analyzing the suppression of cortical AEP as a function of the inter-stimulus interval (Knight et al. 1985; Budd et al. 2013). Parallels between auditory oscillatory responses of rodents and humans have been also reported (Prado-Gutierrez et al. 2012; Venkataraman and Bartlett 2014). In our opinion, the interspecific differences should not be decisive for the mixing results presented here (and previously reported in Prado-Gutierrez et al. 2018) and those obtained by Van Eeckhoutte et al. (2018). Instead of the interspecific differences, we would like to draw attention to the combination of parameters used to estimate the ASSR amplitude in these studies.

### Effect of the analysis parameters on the computation of the ASSR amplitude

The lack of adaptation of the human 40-Hz ASSR reported by Van Eeckhoutte et al. (2018) was obtained using an FFT window of 20.48 s. Such an FFT window is long enough to mask the brainstem ASSR adaptation reported in this study (which occurs in the first 15-30 s of stimulation). Epochs of 20.48 s are two to four times longer than those used in previous studies analyzing the time course of the human 80-Hz ASSR (John and Picton 2000; John et al. 2001; Torres-Fortuny et al. 2011). Remarkably, they are also much longer that those used in basic researches on the human 40-Hz ASSRs, in which the auditory response has been estimated using FFT windows of up to four seconds. These studies include correlation analysis between the 40-Hz ASSR and behavioral thresholds (Van Maanen and Stapells 2005), objective estimates of the loudness growth function based on ASSR (Van Eeckhoutte et al. 2016), the consistency of the 40-Hz ASSR across sessions (McFadden et al. 2014), and the desynchronization of phase-locked neural activities associated with binaural processing (Vercammen et al. 2017).

Results presented in Fig 7 provides an example of how the detection of the ASSR adaptation can be influenced by the analysis parameters. They represent the dynamics of the independent ASSR amplitude, analyzed as a function of the epoch length. More specifically, we computed independent ASSR amplitudes from the original data-set by using different epoch lengths (2.22, 4.45, 8.9 and 17.8 s). Additionally, subsequent epochs were partially overlapped using sliding windows with different sizes. For a given epoch length, we tested 0, 25, 50 and 75% of overlapping. For the different combinations of these parameters, the time evolution of independent ASSRs were fitted to a decreasing exponential function. The adaptation index (*P*_*adapt*_) was calculated for each parameter combination, using the equation:

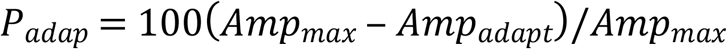

Where *Amp*_*max*_ represents the maximum amplitude of the fitted curve and *Amp*_*adapt*_ represents its asymptotic value.

**Fig 7.**
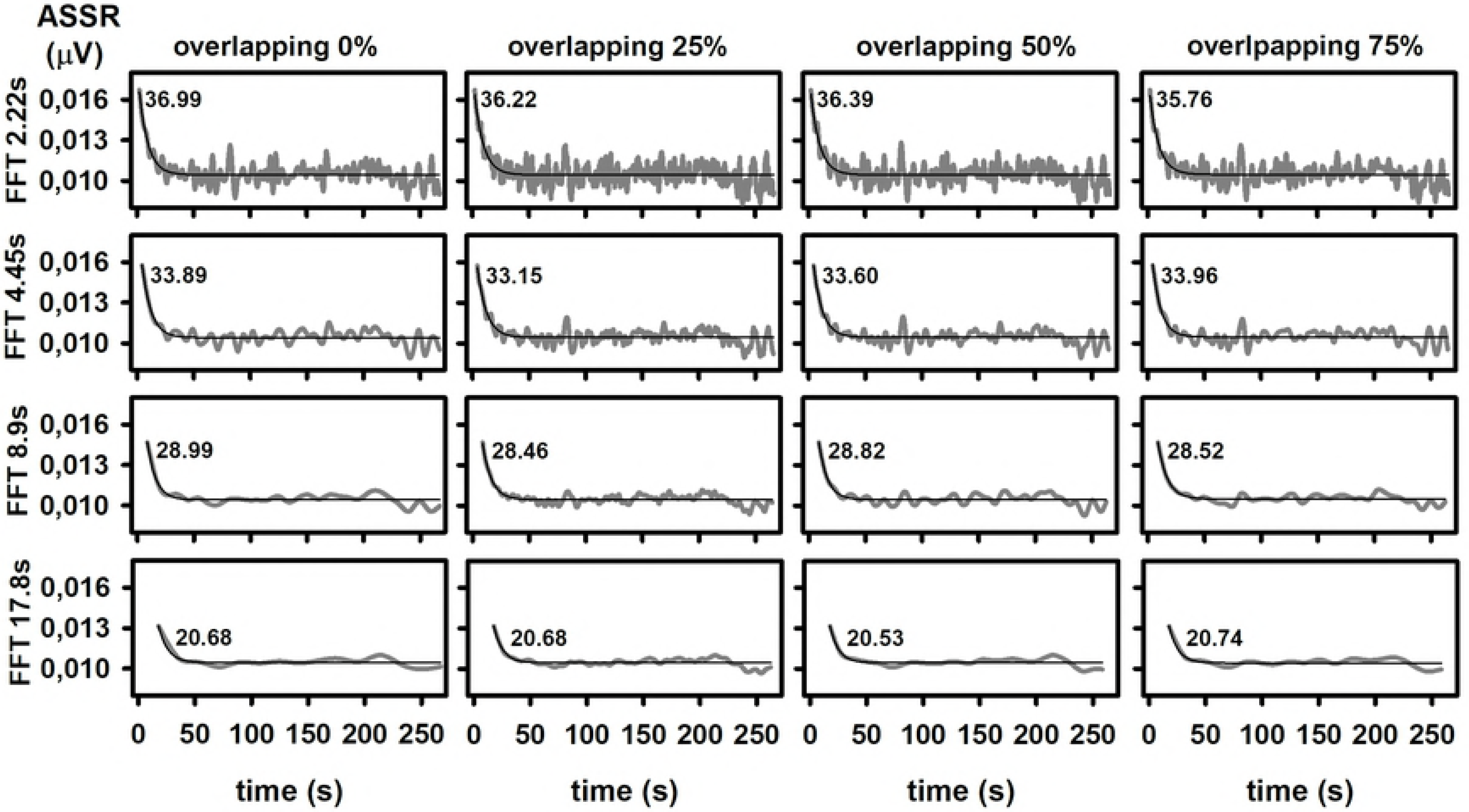
Effect of the epoch’s length (represented in y-label) and epoch’s overlapping on the amplitude of independent ASSR computed by the standard averaging of independent EEG epochs. The dynamics of the ASSR amplitudes (grey lines) were fitted to negative exponential functions (thin black line). Since using different length of FFT windows implies obtaining different number of measurements, ASSR amplitudes are plotted as a function of the recording time. The adaptation indexes are printed inside the corresponding charts.

As shown in Fig 7, the length of the FFT window used for the computation of the response is critical in detecting adaptive behaviors (computed here as the adaptation index). More specifically, the ASSR adaptation is progressively smeared as the length of the FFT window increases. Although greater overlapping implies greater time resolution, such a manipulation did not modify the adaptation index of the ASSR computed with a given FFT window.

### Estimation of dependent ASSR is affected by adaptation

After analyzing the error rate, the detection rate and the recording length of ASSR, Luts et al. (2008) recommend using ASSR detection protocols with a fixed recording length and to judge the significance of the responses at the end of the recording. Furthermore, those authors noted that ASSR can be detected at the initial epochs of a recording, a result that was interpreted as false alarms caused by the greater influence of the background noise when there are few epochs in the averaging. Consequently, they suggest that a minimum of eight epochs should be acquired before evaluating the auditory response (Luts et al. 2008; Choi et al.2011).

Averaging a fixed number of EEG epochs (Fig 3), we corroborated the behavior of the background noise described in previous studies during the averaging of subsequent epochs (John and Picton 2000; John et al. 2001; Luts et al. 2008; Choi et al.2011). More importantly, we demonstrate that the amplitude of the ASSR is higher in the first epochs of the recording and that the dynamics of the dependent ASSR amplitude along the standard and weighted averaging of epochs rely upon the sub-set of epochs selected for the analysis (Figs 3). That behavior is not supported by the dynamics of the RNL, suggesting the ASSR amplitude had been strictly stationary. Therefore, our results indicate that the ASSR amplitudes computed along the sequential averaging of epochs within a recording depends on whether epochs containing unadapted responses are considered or not in the averaging. As mentioned before, the fact of excluding from the averaging EEG epochs with unadapted auditory responses also explains the behavior of the dependent ASSR amplitude associated with sorting averaging.

Consequently, the ASSR amplitude computed at the end of the averaging decreased as more epochs containing unadapted responses are excluded (Fig 4). From a practical point of view, these results highlight the need for defining not just an appropriate length of the recording and averaging stopping criteria for estimating ASSR amplitudes, but also when the response evaluation need to be started.

### Averaging independent vs. dependent epochs

The benefits of estimating the ASSR by averaging independent EEG epochs was evident when analyzing the amplitude and the detection rate of the response (Figs 5 and 6). In practice, the acquisition of independent unadated ASSR could be achieved by replacing the continuous acoustic presentation of tones commonly used to elicit ASSRs by a discrete presentation mode -in which segments of AM-sounds of a few seconds in length are presented after a given inter stimulus interval (ISI). The introduction of the ISI implies that the neural population synchronously responding to the incoming stimulus would be equal or only slightly smaller in size compared to the number of neurons which responded to the preceding stimulation (Presacco 2010; Zacharias et al. 2012). Consequently, the amplitude of the unadated auditory response would remain relatively steady across trials.

Due to the experimental design used in this work, we cannot make any statement about the minimum ISI required for enhancing the amplitude of the ASSR. Future experiments addressing that question are needed. Those studies should focus on the effect of two other aspects of the stimulation strategy: the variability of the ISI and the presentation of broadband noise between consecutive stimuli. As described by Zacharias et al. (2012), a semi-random presentation of acoustic stimuli around a mean ISI might reduce the predictivity of the stimulus, decreasing the magnitude of the response adaptation.

The result presented in Figs 5 and 6 also show that the amplitude and detection of independent ASSR did not vary as a function of the averaging method. In that regard, it is worth noting that our experiments were performed in anaesthetized animals, which were maintained areflexic along the recording session. Therefore, EEG artefacts were extremely uncommon. In this ideal scenario, similar independent ASSR amplitudes are expected to be found when they are computed with the weighted and sorted averaging implemented in this study. However, tools for reducing the effect of EEG artefacts should be determining for the estimation of independent ASSR amplitudes in the clinical practice. The SNR obtained with sorted averaging in higher in comparison with that resulting from weighted averaging (Rahne et al. 2013). This advantage, combined with the fact that sorted averaging does not modify the amplitude of the auditory responses, makes this averaging method in a potential powerful tool to improve the detection of ASSR.

A potential issue regarding the feasibility of the discrete stimulation mode is the possible attenuation of the averaged ASSR amplitudes due to variations in the phase of the neural oscillations from one trial to another. Futures studies need to address this topic experimentally. Nevertheless, the ASSR phase estimated in independent epochs might be less variable than expected, as previous studies have reported a regularity in the expected phase of the human ASSR (Picton et al. 2001, Alaerts et al. 2010; Choi et al. 2011). Similarly, the phase delay -a parameter related to the ASSR latency and that is calculated from the onset phase- have been consistent across studies (Alaerts et al. 2010; Choi et al. 2011). It is worth to note that, even if some phase-related attenuation of the ASSR might be present, our results suggest that applying a discrete stimulation might result in higher ASSR amplitudes than those obtained with the conventional continuous stimulation mode.

In this study, ASSR were evoked by AM-tones with standard sinusoidal envelopes. Modifications of the spectral composition and the amplitude envelops of these standard stimuli have been proposed optimizing the detection of ASSR (e.g. John, et al. 2002; Griskova-Bulanova et al. 2014; Voicikas et al. 2016). Such modifications include the implementation of mixed amplitude and frequency modulated tones (Cohen et al, 1991; John et al, 2001a), AM-tones with exponential envelops (John et al, 2002a, 2004) and AM-noise. Different physiological mechanisms underlie the increase in the ASSR amplitude resulting from the presentation of “modified” acoustic stimuli and that obtained by implementing a discrete stimulation mode. Since it is expected that the ASSR also adapts to the “modified” stimulation, further benefits in the detection of ASSR might be obtained when such stimuli are presented using the discrete stimulation mode proposed here. An ASSR acquisition protocol based on this stimulus paradigm in combination with appropriated averaging methods might lay the foundations for the development of more accurate audiological tests to estimate hearing levels based on ASSRs.

## Acknowledgements

The authors gratefully acknowledge Monica Otero, Universidad Técnica Federico Santa María, Valparaíso, Chile, for the technical support and corrections made to manuscript; Grace A. Whitaker and Wael El-Deredy, Biomedical Engineering School, Universidad de Valparaíso, Valparaíso, Chile, for the corrections made to the manuscript.

## Author Contributions

P.P.G. and E.M.M. have equal contribution to this study. P.P.G. is responsible of the experimental design, data acquisition, data processing and drafting of the manuscript. E.M.M designed the codes for processing the electrophysiological recordings, significantly contributed to the experimental design, data processing and drafting of the manuscript. A.W. contributed to the code design, data processing, and critically reviewed the manuscript. M.Z. critically reviewed the manuscript, making important contributions to its final version.

